# Unexpected global biodiversity responses to climate change arise from complex adaptation, seascape, and dispersal dynamics

**DOI:** 10.64898/2026.03.03.707980

**Authors:** Thomas Keggin, Alexander Skeels, Oskar Hagen, Carlos J. Melián, Conor Waldock

## Abstract

Global marine biodiversity peaks in the tropics and is widely projected to shift polewards under climate warming as species track thermal optima. These projections rely largely on phenomenological models that treat species responses as shifts in climate suitability and neglect fundamental eco-evolutionary dynamics. The consequences of these dynamics in fluctuating populations for global biodiversity patterns therefore remain unresolved. We addressed this deficit by integrating local adaptation, dispersal, and gene flow – mediated trait homogenisation into spatially explicit global simulations of >1,400 tropical reef fishes under climate change. Biodiversity responses departed systematically from simple poleward redistribution. Depending on the interaction between dispersal and adaptive capacity, latitudinal richness peaks widened or narrowed, shifted equatorward or poleward, or collapsed entirely. These alternative regimes emerged from threshold-dependent interactions between seascape connectivity, metapopulation trait homogenisation, and accumulating thermal stress. Critically, greater dispersal enhanced persistence only below seascape-driven connectivity thresholds; beyond which homogenisation of traits across thermally heterogeneous populations disrupted local adaptation – particularly at metapopulation peripheries – and increased extinction risk. Continuous demographic feedback further generated cumulative stress and extinction tipping dynamics absent from snapshot suitability models. Our results show that eco-evolutionary dynamics can invert or destabilise expected climate-driven redistribution patterns, indicating that projections excluding these mechanisms may systematically misestimate the magnitude and direction of global biodiversity change in a rapidly warming ocean.

## Introduction

Rapid ocean warming is already redistributing marine biodiversity worldwide, largely determined by populations’ ability to track shifting thermal niches and their ability to persist via adaptation (Figure 1) – with extinction risk rising where both responses fall short (Downie et al., 2025; Pecl et al., 2017). It follows that species with higher dispersal abilities and greater capacities for local adaptation are expected to be least vulnerable to climate change (Feary et al., 2014). Considering both dispersal and adaptation, the benefits of each could be additive, with species that are both highly mobile and adaptive persisting best within a dynamic environment. However, dispersal between geographically isolated populations can result in gene flow and subsequent homogenisation of divergent phenotypes across a species’ range (Capblancq et al., 2020; Lenormand, 2002; Slatkin, 1987). If connected populations are locally adapted to different environmental conditions, phenotypic homogenisation across a metapopulation may introduce divergent phenotypes with a better or worse fit to a population’s local environment – exacerbated for populations on the geographic or environmental periphery (Lenormand, 2002). This inter-population genetic exchange can therefore disrupt local adaptation (Crispo, 2008; Nosil, 2009; Slatkin, 1987). Whilst the emergent consequences of a dispersal-adaptive dynamic have been explored at smaller scales for single or few species (Aguirre-Liguori et al., 2021), it is unclear how the dynamics of adaptation and dispersal modify emergent patterns in large-scale biodiversity responses to climate change (DeMarche et al., 2019).

**Figure 1:**
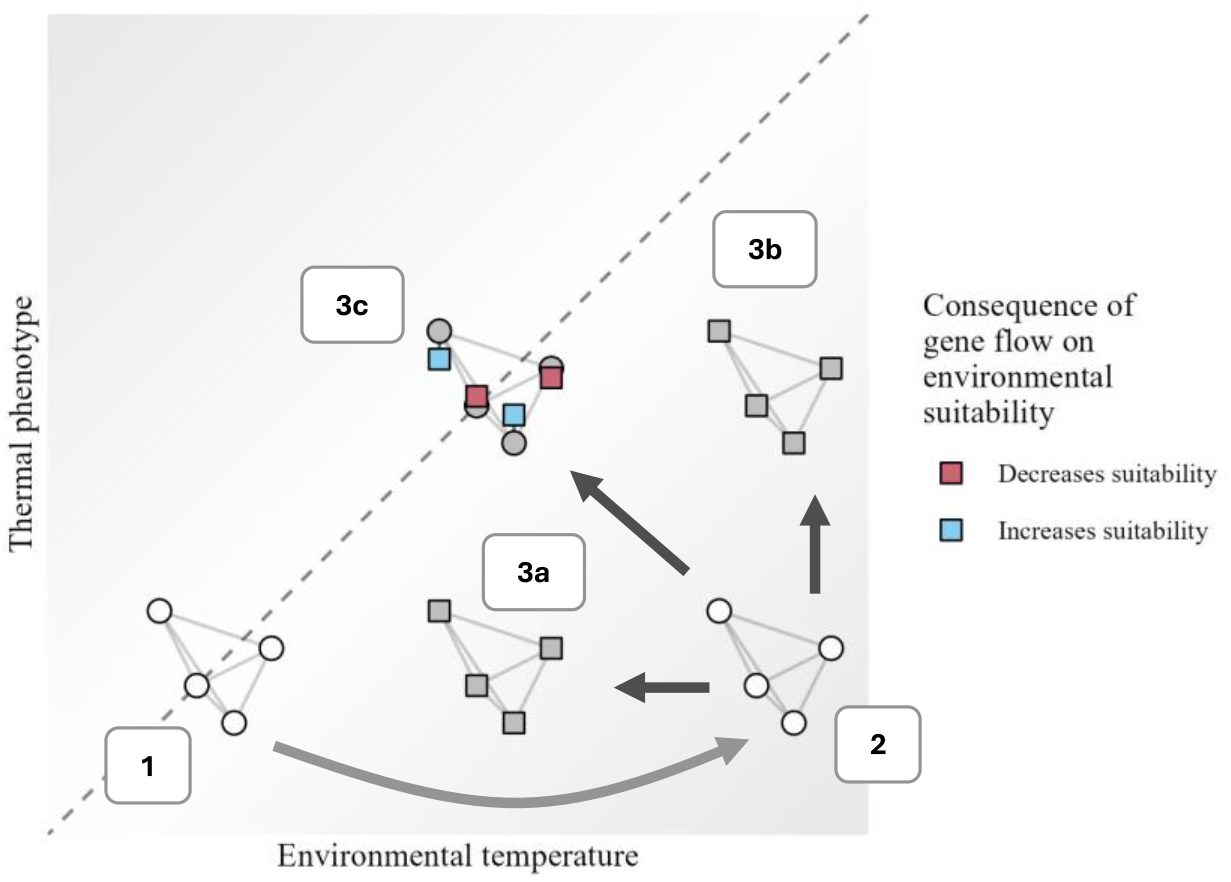
Pathways for the response of metapopulations to environmental change through adaptation, dispersal, and gene flow. The grey dotted line represents perfect environmental suitability, the light grey arrow represents the environmental change, and the dark arrows represent metapopulation responses. Circles indicate populations with no, or incomplete, response; squares indicate populations after a complete response. 1. Environmental suitability at *t* – 1. 2. No tracking mechanism. The environmental temperature increases, but the populations do not respond, leading to decreased suitability. 3. Species respond to changing environment. 3a. The environmental temperature increases locally, but populations can move towards suitable habitat – changing the environment relative to them and “moving back” through environmental space. 3b. The environmental temperature increases and the populations cannot track their niche space, instead their thermal niche tracks the environment through local adaptation. 3c. The environmental temperature increases and the populations respond by both moving to suitable habitat and adapting. However, with trait homogenisation across the metapopulation through gene flow, the efficacy of adaptation is disrupted.

The latitudinal diversity gradient of increasing species richness (LDG) from the poles to the equator is a first-order global biodiversity pattern (Tittensor et al., 2010; Willig et al., 2003) that is already being disrupted by climate warming (Edgar et al., 2023; González-Barrios et al., 2025). Tropical marine reef-associated fishes follow this latitudinal species richness gradient, with the contemporary richness pattern being associated with habitat availability and environmental temperature patterns through both deep (Gaboriau et al., 2019; Mannion, 2020) and shallow time (Tittensor et al., 2010). Their spatial distributions and abundances also show a strong response to thermal gradients (Auber et al., 2022; Tittensor et al., 2010), with tropical species already living near their thermal limits (Vaughan et al., 2025; Waldock et al., 2019), leading to tropical populations being particularly vulnerable to climate warming (Chaudhary et al., 2016; Edgar et al., 2023). Climate warming is therefore expected to disrupt and shift the overall latitudinal diversity gradient in species richness as species track suitable thermal habitat towards the poles (Auber et al., 2022; Munday et al., 2008).

These predicted polewards range shifts, however, assume that species responses are independent of local adaptation to increasing temperatures, that dispersal ability is largely unlimited across regional to global seascapes, and that no interaction exists between dispersal and local adaptation (A. Lee-Yaw et al., 2022; Aguirre-Liguori et al., 2021; Bayliss et al., 2022; Urban et al., 2022). Yet in reef fishes, as in many other classic patchy metapopulation systems (Hanski, 1994), dispersal occurs across geographically and thermally complex seascapes of semi-discrete reef patches through time (Mora, 2015). Characteristics of these seascapes determine thresholds in species responses to environmental change (Monaco & Helmuth, 2011) and the effective distances (effective relative to species’ active or passive dispersal behaviours) between reef patches form barriers to dispersal across a seascape (Mora, 2015). Similarly, with a warming climate, limits to adaptive capacity relative to local environmental change can determine whether species can locally persist – *i*.*e*., if environmental conditions change faster than species’ adaptive capacities, they likely perish (Lavender et al., 2021). Expectations for both dispersal and thermal thresholds have been explored for global species responses to climate change in both static and dynamic seascape structures (Keggin et al., 2023; Munday et al., 2008). However, how these global thresholds will be modulated by the interaction amongst adaptation, dispersal mediated gene flow on metapopulations, and dynamic seascapes remains unexplored. This is especially pertinent considering the large variation in species’ dispersal abilities (Green et al., 2015) and the complexity of adaptive responses (Tigano & Friesen, 2016).

The impact of known patch-to-patch dispersal limitations, variable local adaptive capacity, and the disruptive impact of gene flow on phenotypes on global scale biodiversity responses remains largely unknown (Alzate & Hagen, 2024) – even whilst we actively monitor the impacts of a warming climate on global biodiversity patterns.

Spatially explicit eco-evolutionary simulations can incorporate dispersal, local adaptation, and dispersal mediated gene-flow across populations – and allow these processes to emerge into global biodiversity responses to environmental change (Hagen et al., 2021; Haller & Messer, 2023; Urban et al., 2022). Simulations have been widely applied at the individual level and often unveil different biodiversity responses to environmental change when compared to multivariate associative, or pattern driven, predictions at small spatial scales (Baiotto & Guzman, 2025; Benning et al., 2023, 2024; Booker, 2024; Melián et al., 2011; Vahdati & Melián, 2022). For example, dispersal simulations validated against genomic and environmental data show that marine rafting crustaceans experiencing gene-flow between dispersing populations could be either reducing or exacerbating maladaptation to changing environmental conditions – impacting their range shifts with climate change (Liu et al., 2026). Under the same dynamics, alpine species may persist longer in habitats than expected when local adaptation is incorporated in population responses, yet such persistence comes at a cost for species with highly connected metapopulations as traits track to the populations’ average values – which can increase local maladaptation (Cotto et al., 2017; DeMarche et al., 2019; Moore et al., 2007). Recent developments in population-level eco-evolutionary simulations allow us to scale the simulation approach from few populations in small areas to thousands of species across global geographies (DeMarche et al., 2019; Hagen et al., 2021; Urban et al., 2022). We are now able to explore how global latitudinal shifts in tropical reef fish species diversity might be modulated by varying the seascape, warming, and dispersal dynamics at the local, regional, and global scales.

We configured global, forward-in-time simulations under projected future climate change scenarios – incorporating implementations of local adaptation, dispersal, and the trait homogenisation consequence of gene flow to understand their contributions to global biodiversity responses. We assumed a dispersal, mating, then selection paradigm (see methods) that follows the gen3sis engine’s internal framework (Hagen et al., 2021; Lakovic et al., 2017). We investigated the responses of 1,427 tropical reef fish species between 2014 and 2100 under the CMIP6 RCP8.5 scenario, using contemporary distributions, abundances, and phylogenetic information (Auber et al., 2022; Rabosky et al., 2018) across the global extent of shallow-water reef habitats. For all cells within all species across annual time steps, we simulated population-level dispersal events between habitat patches with associated trait homogenisation, changes in population distributions of local thermal optima, and logistic population growth modulated by thermal suitability. We measured emergent population-to-species responses to changing sea surface temperature (SST). We tracked gradients in species richness, temporal compositional turnover, species’ range size and range fragmentation, and changes in the spatial pattern of species distributions. By varying the adaptive and dispersal capacities across simulations, we tested how dispersal with trait mixing, local adaptation, and their interaction might impact these response metrics; and ultimately how incorporating additional eco-evolutionary mechanisms will fill gaps in our understanding of how global biodiversity responds to climate change.

## Results and discussion

### Identifying thresholds in species responses to climate warming

Across different simulated dispersal ranges and adaptive rates, we found substantial variation in biodiversity response patterns by the year 2100. Our simulations could be best aggregated into six unsupervised groups based on global biodiversity metrics (Fig. 2, Supplementary Fig. 1). Whilst some groups aligned with existing expectations from associative models, others, belonging to specific parts of the adaptation-dispersal gradient, did not (Fig. 2). These groups were defined by global-species richness, -mean cell species richness, -mean species abundance, -mean range size (Fig. 2a,b), -reef occupancy, and -mean meta-population connectivity at the year 2100 (Supplementary Fig. 1). These six response groups reflected spatial responses in the latitudinal species richness gradient (Fig. 2c), which mostly formed a Gaussian latitudinal gradient in each equator (Supplementary Fig. 2), also in line with previous works (Auber et al., 2022; Munday et al., 2008). Beyond previous expectations, however, we found both poleward and equatorward movement of richness peaks between 2014 and 2100 (Fig 2c; Supplementary Fig. 3). The shape of latitudinal richness peaks was also variable depending on dispersal and adaptation parameters with peaks becoming narrower (Fig. 2: gene flow and adaptation interplay, and adaptation limitation groups), wider (Fig. 2: species expansion group), flatter (Fig. 2: dispersal limitation) or breaking down entirely (Fig. 2: species collapse group) – with some instances of global extinction (Fig. 2: global extinction group). When both dispersal ability and adaptive rate values were low, simulations experienced high levels of extinction (Figure 2a: Global species richness), with five simulations experiencing global extinction by 2100. Simulations with high adaptive rate and low dispersal had narrower latitudinal richness peaks moving towards the equator (Supplementary Fig 3). Where adaptive rate was limited and dispersal values were high, simulations experienced a narrowing of richness peaks as they moved polewards, in line with broad expectations when assuming largely unrestricted dispersal and static thermal niches – as in species distribution models (Auber et al., 2022). This variation, split across six distinct groups, expands on simpler patterns reported in associative global models, *i*.*e*., that richness will shift polewards with climate warming (Munday et al., 2008).

**Figure 2:**
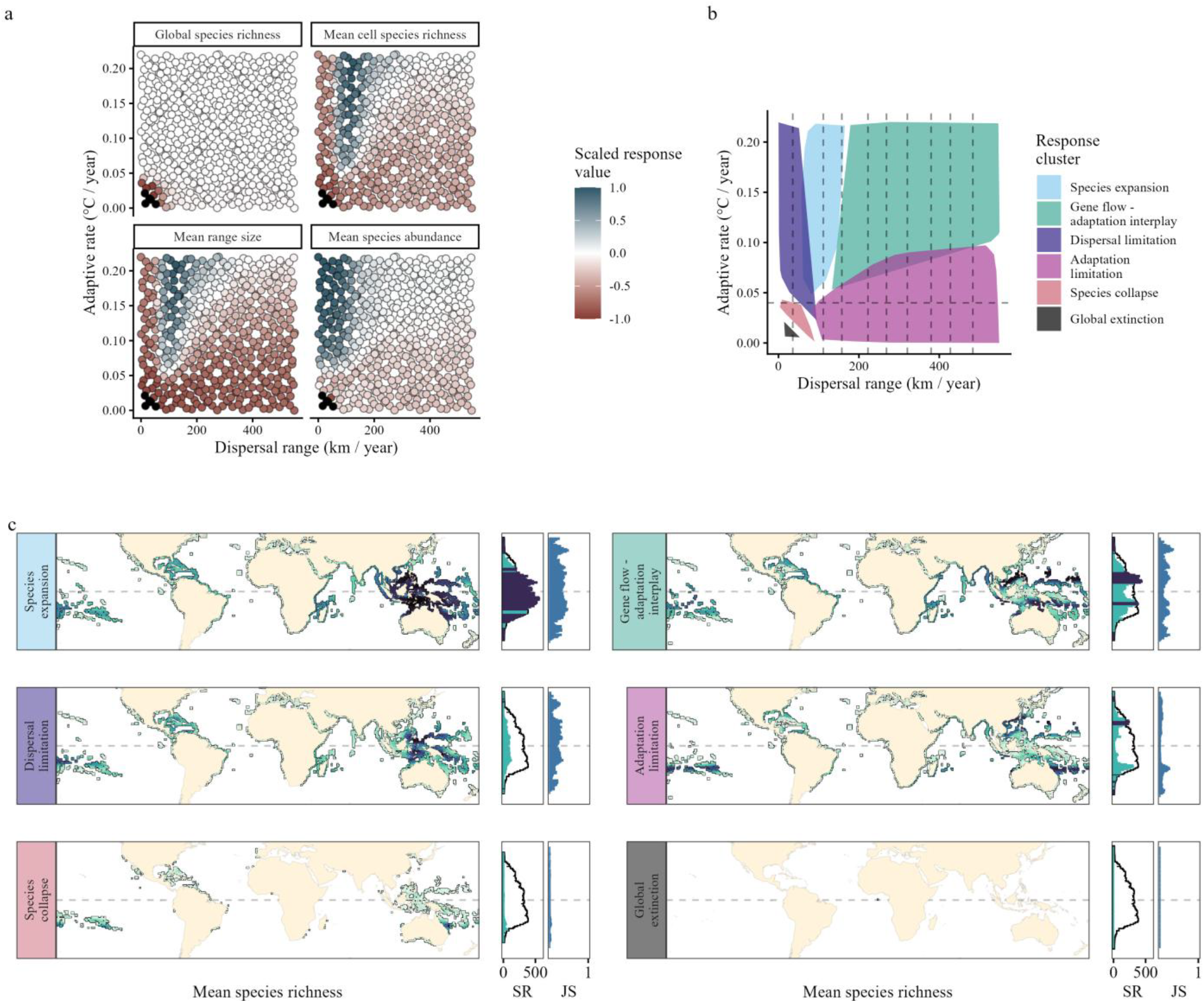
Globally aggregated species responses to sea surface temperature change between the years 2014 and 2100 across adaptive rate and dispersal range gradients. a) Global response metrics of simulations (points) with colours indicating the difference between 2014 and 2100 values. Values are scaled from -1 to 1 where 0 is no change and +/-1 is the maximum absolute change across simulations. Blue values indicate an increase in response metric through time, and red a decrease. Black indicates global extinction by 2100. b) Simulations were best clustered into six groups based on global response metrics. Each coloured polygon envelopes a response group. The dashed horizontal and vertical lines respectively represent thermal and dispersal thresholds in the global seascape (Supplementary Fig. 4). c) Panels representing mean aggregated species richness and compositional turnover. Maps represent species richness at the year 2100 with darker colours indicating greater richness. The SR panel is mean latitudinal species richness where the black line is species richness at 2013, and the columns are species richness at 2100. Dark and light columns indicate an increase or decrease in species richness, respectively. The JS panels represent mean latitudinal species compositional turnover between 2013 and 2100 as measured by Jaccard’s similarity index. Values of 1 indicate identical species composition and 0 indicates complete turnover.

The boundaries between the six response groups in parameter space aligned with environmental thresholds in the seascape (Fig. 2b, Supplementary Fig. 4). We identified a single thermal threshold: the mean annual temperature increase across all habitable cells and years in the simulated seascape. Positive responses with warming (increased species richness, greater range size, higher abundances) were only realised if the adaptive rate parameter value was greater than the mean global annual increase in SST between the years 2014 and 2100 (figure 2b, horizontal dashed line). Below this mean global annual SST increase, simulations grouped into dispersal dominated dynamics, species collapse, or even global extinction – similar to species collapse and extinction responses in shallow water marine systems during previous hyperthermal events during the Meso- and Cenozoic (Dunhill, Foster, Azaele, et al., 2018; Dunhill, Foster, Sciberras, et al., 2018; Kiessling & Aberhan, 2007; Salles et al., 2025). These declines were only rescued, non-linearly, by very high values of dispersal (*e*.*g*., in the adaptation limitation group). On the other hand, when the adaptive rate was greater than mean annual SST increase, simulations were either limited by dispersal, responded positively to global SST change, or experienced a more complex non-linear response in parameter space – all dependent on dispersal ability.

Geographic thresholds in the seascape were identified as sharp peaks in the distribution of habitable cell-to-cell distances (Supplementary Fig 4a). As expected, dispersal abilities below the minimum distance between habitable reef cells, in tandem with insufficient compensating adaptive rate, resulted in abundance declines, range contractions, range fragmentation, and extinctions. Overcoming this initial barrier allowed species to reach their most expansive response values in species richness (highest), range size (largest), and global reef occupancy (greatest occupation; Fig. 2a, Supplementary Fig. 1,4d) enabling species to track suitable habitat and to adapt to the warming environment. The benefit of dispersal was greatest when dispersal ability matched the second peak in cell-to-cell distances of 111.5 km (Fig. 2, Supplementary Fig. 4d). After this second peak, many response values were negatively disrupted by trait homogenisation within metapopulation clusters and decreased non-linearly dependent on both dispersal ability and adaptive rate. Whilst the specific threshold value emerges from the discretised seascape, its magnitude overlaps with effective dispersal estimates for reef fishes (25 – 150 km (Palumbi, 2003)), suggesting that effective connectivity relevant in real marine systems may plausibly align with the hypothetical dynamics explored here. And whilst these sharp dispersal thresholds, along with the thermal adaptive rate threshold, are likely less prominent in the real world – depending on habitat patch structure and the continuity of populations (Berkström et al., 2020; Kulbicki et al., 2014; Selkoe et al., 2008, 2016) – they have been found to occur in experimental metapopulations (Molofsky & Ferdy, 2005). Regardless, they provide critical geographic context to the interaction between adaptation and dispersal through trait homogenisation dominant in our simulations. Through gaining understanding of these *in-silico* scenarios, we can form expectations of how biological processes could interact with geographic constraints across real seascapes to set the context of global species responses to ongoing climate change.

### The complex interplay between local adaptation and dispersal with trait homogenisation

Our simulations showed an important interaction effect between population-level dispersal and local adaptive rate in the emergence of species responses to climate change – one that is modified by thresholds in reef habitat structure and thermal dynamics through time. In real systems, the extent of trait homogenisation through gene flow is highly variable and dependent on the selective, genomic, and behavioural context (Fitzpatrick et al., 2015; Raeymaekers et al., 2014; Tigano & Friesen, 2016); but even through abstraction, our simulations added significant complexity to emergent species responses to climate change beyond what existing phenomenological models predict (Fig. 2) (Auber et al., 2022; Munday et al., 2008). To understand how the interaction between dispersal and local adaptation modulated these responses, we isolated the trait homogenisation function over a single time step. By holding the adaptive rate, range size, and environmental heterogeneity constant, we can see the impact of the adaptation-dispersal interaction on environmental suitability across metapopulations with differing dispersal abilities. Trait homogenisation within metapopulations had a variable consequence on environmental suitability across constituent populations dependent on their connectivity through dispersal ability (Fig. 3). Under high dispersal, more thermally distant cells were connected. This led to homogenisation across more disparate trait values, and therefore greater absolute trait change and in turn to more instances of maladaptation – whereby mean population thermal trait values were more mismatched with the environment than they were initially. This increase in maladaptation was reduced towards the mean metapopulation trait space, with populations closely trailing the mean experiencing beneficial gene flow as their trait values were “pulled” towards the local SST value by connected warmer adapted populations (Fig. 3, Supplementary Fig. 5). Conversely, populations on the thermal peripheries, *i*.*e*., most distant from the metapopulation mean trait value, experienced the greatest levels of maladaptation as their thermal traits were disrupted by more disparate trait values through trait homogenisation, leading to more local extinction events and to range contractions. Although the periphery effect dominated our model configuration and is well documented geographically as a constraint of gene flow on local adaptation (Lenormand, 2002), its large-scale influence remains poorly explored. Next, we show how meta-population periphery effects can strongly modify global biodiversity responses.

**Figure 3:**
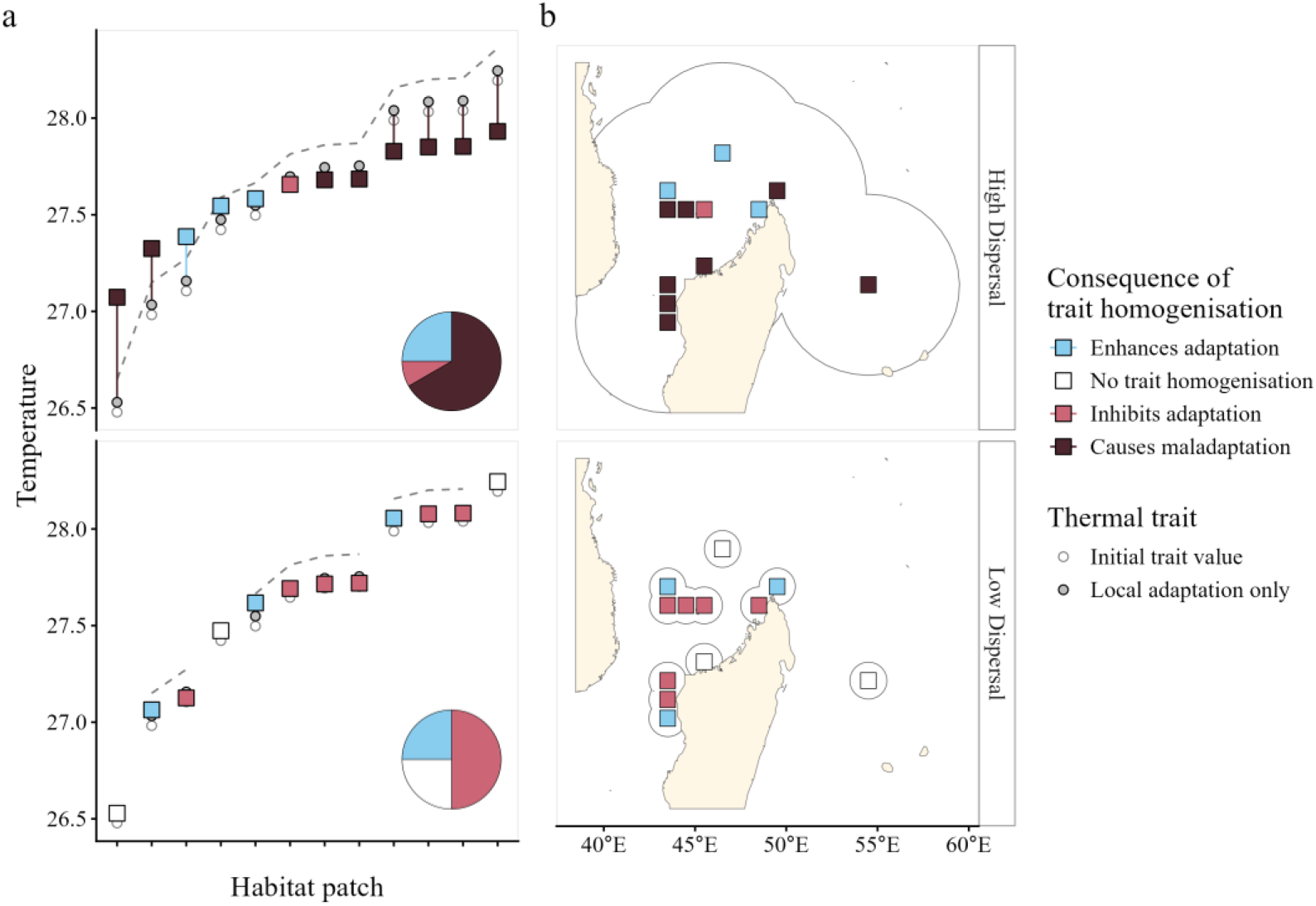
The impact of trait homogenisation (without distance decay) on metapopulation adaption under high and low dispersal regimes (top and bottom panel rows, respectively). Colours indicate the consequence of trait homogenisation: dark red indicates maladaptation; light red indicates inhibition of adaptation; white indicates no effect; and blue indicates enhancement of adaptation. a) Adaptive outcomes across habitat patches. The grey dotted line represents sea surface temperature. Line continuity reflects metapopulation connectivity, as represented by the map polygons in (b). Clear circles indicate the initial phenotype (thermal trait at *t*-1), grey circles represent the phenotype after the first half of the evolution function has been applied (local directional adaptation only), and squares indicate the phenotype after the second half of the evolution function has been applied (abundance-weighted trait homogenisation across metapopulations). Square colours indicate the fitness outcome of the phenotype homogenisation step. b) Spatial structure of the habitat patches corresponding to (a), illustrating the effects of gene flow. Transparent polygons indicate the dispersal range from each patch; patches within a contiguous polygon form a panmictic metapopulation.

The emergent consequence of the dominant metapopulation periphery effect explained variation in global biodiversity responses to climate change. Across simulations, phylogenetic mixed-effects models showed that populations consistently grew less suited to their environment as they approached the thermal periphery, or edge, of their metapopulation (*P* < 0.001; positive slope estimates, Supplementary Fig 5a). Subsequently, peripheral populations suffered reduced environmental suitability (*P* < 0.001), which reduced local abundance (*P* < 0.001) and led to local extinction and truncated range sizes. We found that the magnitude of this thermal suitability decrease towards the periphery also differed across ecoregions in our seascape (Supplementary Fig. 5b) (Spalding et al., 2007). These differences could be explained by the magnitude of thermal increase over time. Ecoregions with a greater thermal increase showed a reduced periphery effect (e.g., Northern and Central Red Sea, Saharan Upwelling, Leeuwin, and Northern Bay of Bengal), due to the balancing of trailing and leading populations when all populations are thermally adapting in the same direction (*i*.*e*., to warmer conditions). Similarly, ecoregions with relatively sparser habitat patches showed a marginally significant, reduced periphery effect (*P* = 0.09, *e*.*g*., Cape Verde, Tonga Islands, and South New Zealand) – equivalent to decreased metapopulation connectivity at low dispersal capacities (Figure 3; Supplementary Fig. 5) (Slatkin, 1987). Given that geographic and thermal distances between cells are positively correlated for every year (linear regressions, 0.04 < Adj. R^2^ < 0.05, *P* < 0.001, Supplementary Fig. 6), increasing metapopulation size through higher dispersal also increases experienced, or apparent, environmental heterogeneity. The heterogeneity across the metapopulation in turn leads to more disruptive trait homogenisation, reduced periphery population abundance and central population dominance (Lenormand, 2002). At the global scale, the emergent result in our simulations is a narrowing of species ranges and latitudinal richness peaks when dispersal abilities are higher than what can be mitigated against by local adaptive processes.

### Latitudinal diversity gradient collapse dynamics

The loss of a Gaussian latitudinal species richness response pattern with climate warming was correlated with species declines and included high levels of extinction (Fig. 4, Supplementary Fig. 2,3). This correlation between climate warming-driven species declines and the collapse of a definable LDG pattern is not without precedent – it reflects historic evidence of the flattening of the latitudinal diversity gradient during the hyperthermal Permian-Triassic mass extinction event (Rummer & Munday, 2017; Song et al., 2020). Whilst these outcomes were extreme, systematic exploration of the full parameter space revealed mechanisms and conditions which can lead to mass extinctions. Simulations with the least ability to respond to climate change (low dispersal and adaptive rate) were the first to experience species declines and extinctions (Fig. 4c,d). We found that abundance and distribution declines in response to thermal stress compounded from one year to the next (Figure 4d), resulting in faster species declines than expected through the thermal suitability of reef cells at each time step alone (Figure 4a,b). Species declines and subsequent extinctions were best described using a multiyear predictive modelling framework (Supplementary Fig. 7), where declines could be predicted by cumulative thermal stress over one to several years and extinctions could be predicted by cumulative thermal stress over four or more years (Supplementary Fig. 7). These extinctions were in turn dominated by tipping points in the thermally dynamic seascape, where inverse exponential declines were triggered sooner, and played out faster, in poorly responding simulations (Fig. 4). By capturing species decline dynamics through cumulative stress, global levels of biodiversity patterns such as the latitudinal diversity gradient and extinction events might occur faster than expected by modelling approaches based on commonly implemented temporal point estimates of environmental suitability through time. This approach highlights the potential to investigate known cumulative thermal stress and signals of abundance change as an early warning signal of population extirpation and species loss at the global scale (Baruah et al., 2020).

**Figure 4:**
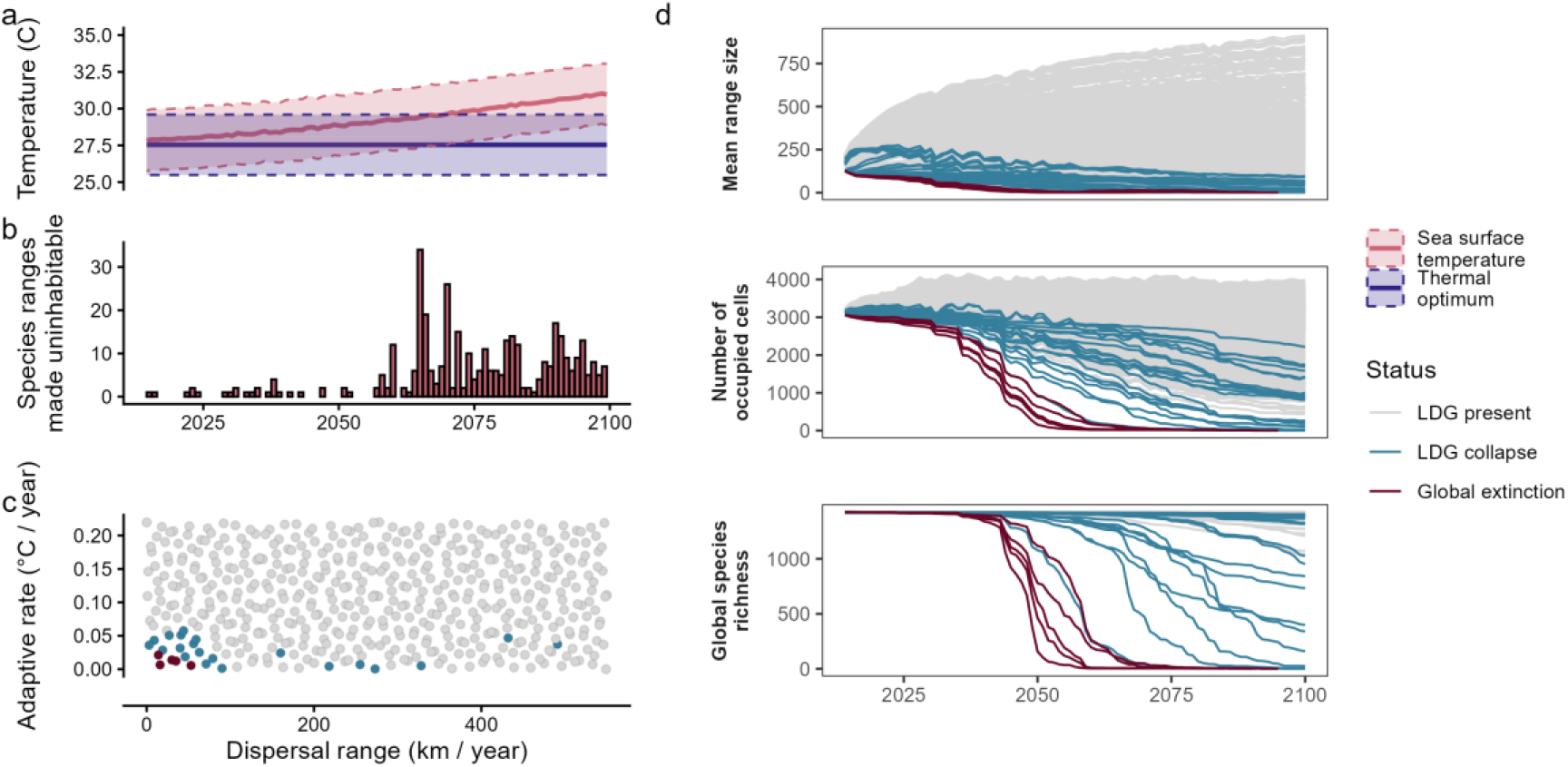
Timeseries of latitudinal diversity gradient collapse. a) Thermal optima across species vs the sea surface temperature of the cells they occupy through time. Thick blue and red lines represent the mean thermal optimum across species on simulation initialisation and SST of the inhabited cells, respectively. The shaded ribbons represent thermal optima and SST values within 1 standard deviation of the mean. b) Given maximum starting abundance, the number of species whose entire range becomes uninhabitable each year through thermal stress, given no dispersal or local adaptation. c) Simulation parameter values highlighting the simulations in which latitudinal diversity gradient patterns collapse (blue), or where there is global extinction (dark red). d) Timeseries for each simulation showing how species richness and underlying vulnerability metrics change through time, highlighting latitudinal diversity gradient collapse simulations (blue), and global extinction simulations (red).

### Comparison with current associative methods

Our results align with decades of work exploring how gene flow modifies trait adaptation and population dynamics, but find that introducing these mechanisms to global biodiversity-climate response modelling generates differing response patterns. Species richness patterns derived from a species distribution modelling framework (without local adaptation or trait homogenisation) using the same species subset (Auber et al., 2022) most closely align with simulations containing an adaptive rate above 0.04 C° / year (higher than the mean annual global SST increase) and dispersal values around 111.5 km / year (first dispersal distance peak; Supplementary Fig. 4). This corresponds to our “best-case” scenario grouping of “species expansion” whereby richness is retained in the tropics and expands equator-wards (Fig. 2, Supplementary Fig. 3,8). Counterintuitively, this low dispersal and high adaptive parameter combination is the opposite of the implicit assumptions of associative modelling frameworks which assume fixed thermal phenotypes and only regional restrictions to dispersal. By outputting similar patterns through different assumed processes, we highlight how introducing eco-evolutionary mechanisms into global thinking can disrupt our understanding of how species respond to rapid environmental change.

## Conclusions and future directions

Eco-evolutionary theory has long emphasised the role of gene flow in shaping adaptation, yet the influence of this crucial interaction remains largely overlooked and unexplored in large-scale temporo-spatial contexts modelling biodiversity responses to climate change. Our findings bridge this gap, revealing that dispersal dynamics, when coupled with adaptive processes, produce a range of potential global responses that could significantly impact expectations of how species might shift and experience extinction risk, and how the existing latitudinal diversity gradient in species richness might be altered. The complexity of these interactions, highlighted by phenomena like the periphery effect, suggests that current models may vastly under- or over-estimate the resilience of species in the face of a warming environment. We find that by including even simplistic adaptive processes and their interaction with dispersal and trait homogenisation across metapopulations underscores the need for us to re-evaluate our expectations for global species responses to climate change.

## Acknowledgements

We would like to thank: Sarah Schmid for feedback and sound boarding; Bruno Mäser for computational support; Jakob Brodersen and Philine Fuelner for helpful feedback on abstraction terminology; the Computational Phylogenetic Group, UNIL, for important early constructive criticism on conceptualisation of mechanisms; Theo Gaboriau for conceptual discussion of population level simulation frameworks; Nadja Pepe, Michelle Achermann-Sidler, and Patricia Achleitner for patient administrative support; and Dominik Kirschner, Anna Kätzlinger, Yohann Chauvier, and Lena Roux for “invisible”, kind personal support.

## Supplementary Figures

**Supplementary figure 1:**
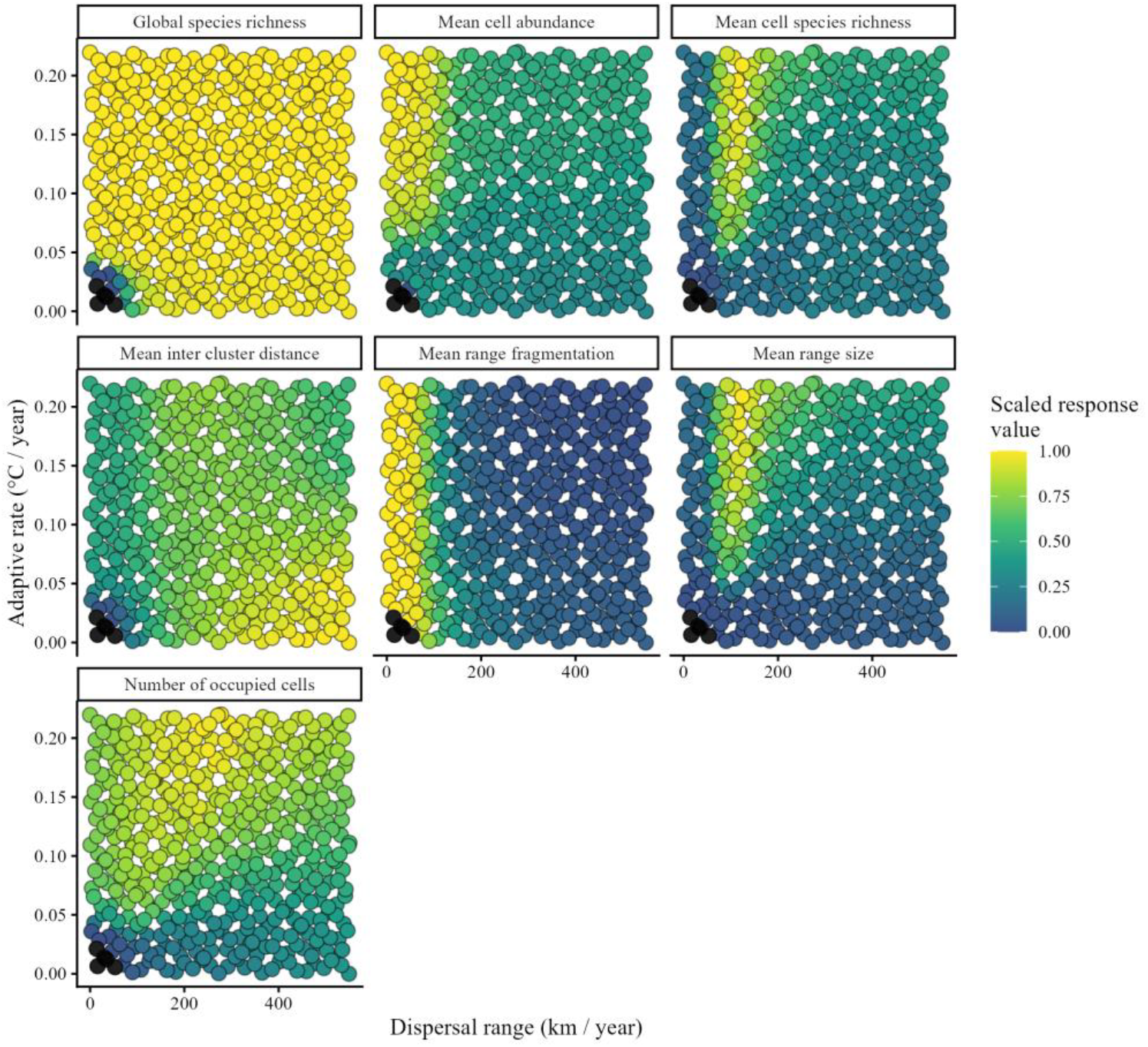
Scaled global response metric values at 2100 in dispersal and adaptive parameter space. Each panel is a response metric. Each filled point is a simulation. Fill values are global response metric values, scaled from 0 (darkest and lowest) to 1 (lightest and highest). Solid black filled points are simulations that have gone globally extinct by 2100.

**Supplementary Figure 2:**
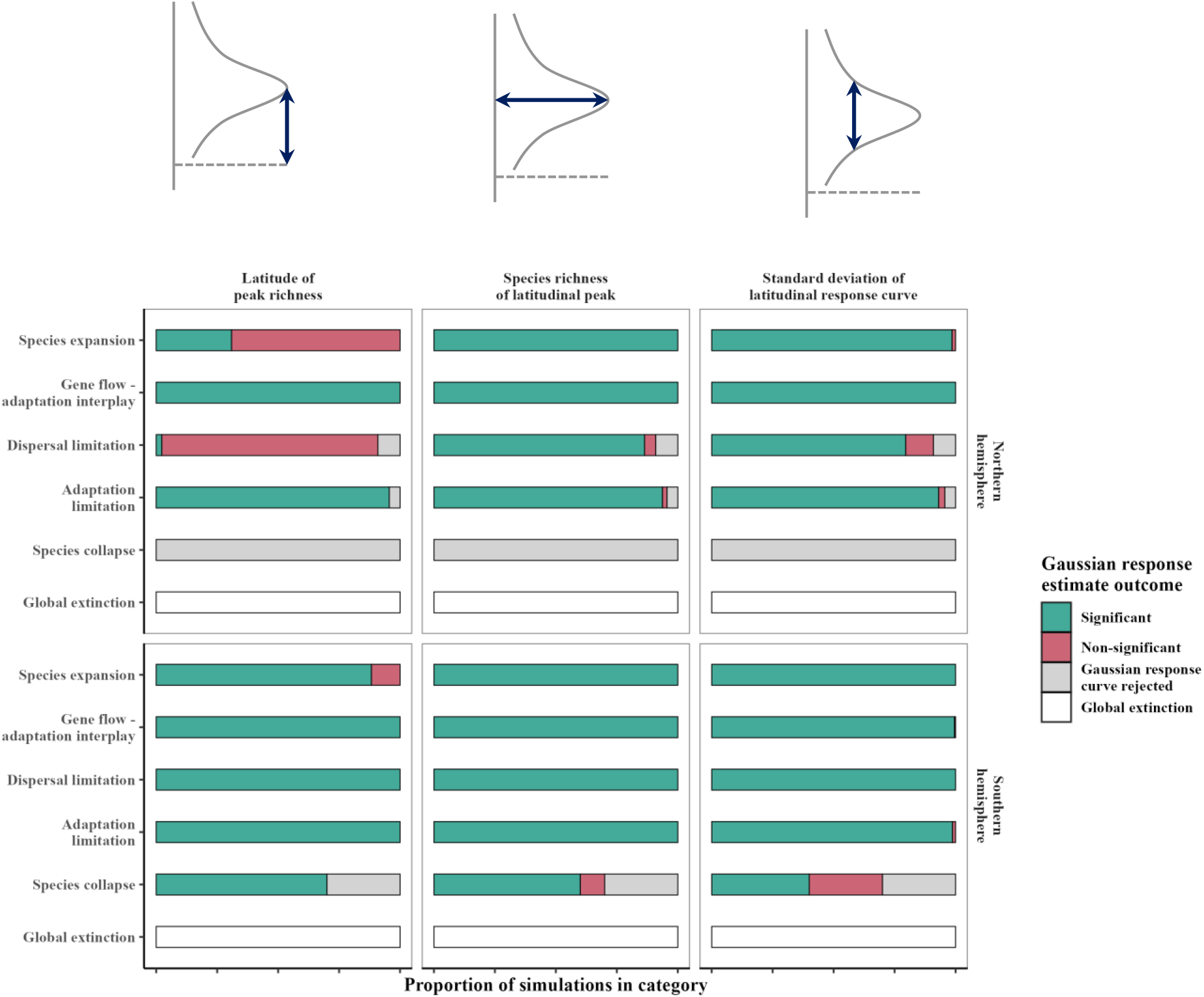
The proportional suitability of a gaussian latitudinal diversity gradient across simulations grouped by their global responses (Fig. 2). Each bar represents the proportional divide of simulation groups into significant Gaussian distribution estimate fit, non-significant Gaussian distribution fit, Gaussian fit rejected in favour of greater support for a uniform distribution, and global extinction – in each hemisphere.

**Supplementary Figure 3:**
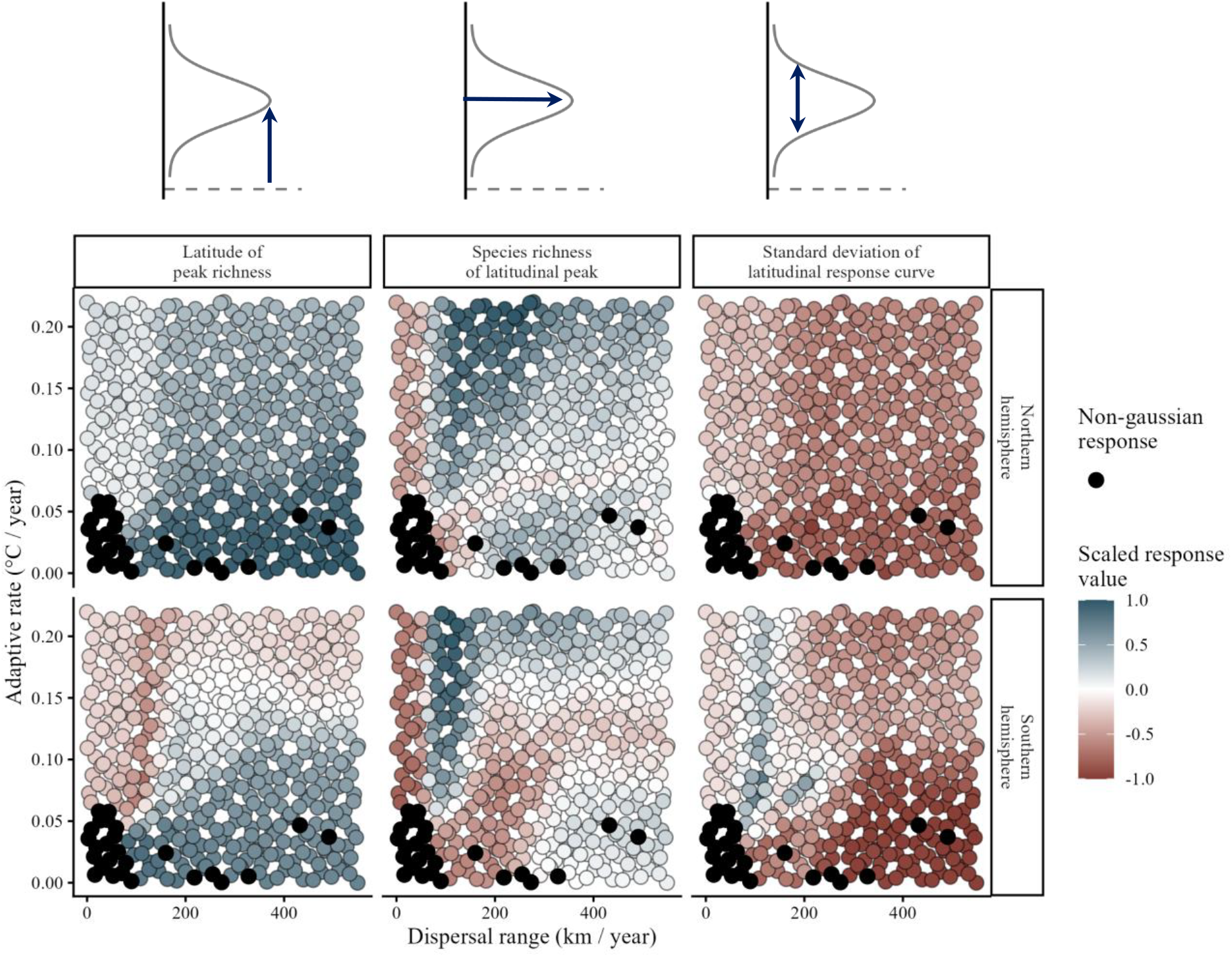
The latitudinal diversity gradient response for simulations that best fit a bimodal Gaussian distribution (Gaussian in both hemispheres). The distribution in each hemisphere is characterised by the richness peak latitude (mean), richness peak value (species richness), and the richness peak width (standard deviation). Blue denotes an increased metric value in 2100 compared to 2014, and red a decrease. White fill indicates no change. Black fill indicates where the latitudinal diversity gradient could not be described by a gaussian response (either extinct, a non-significant fit, or the response was better supported for a uniform distribution – see Supplementary Figure 2).

**Supplementary Figure 4:**
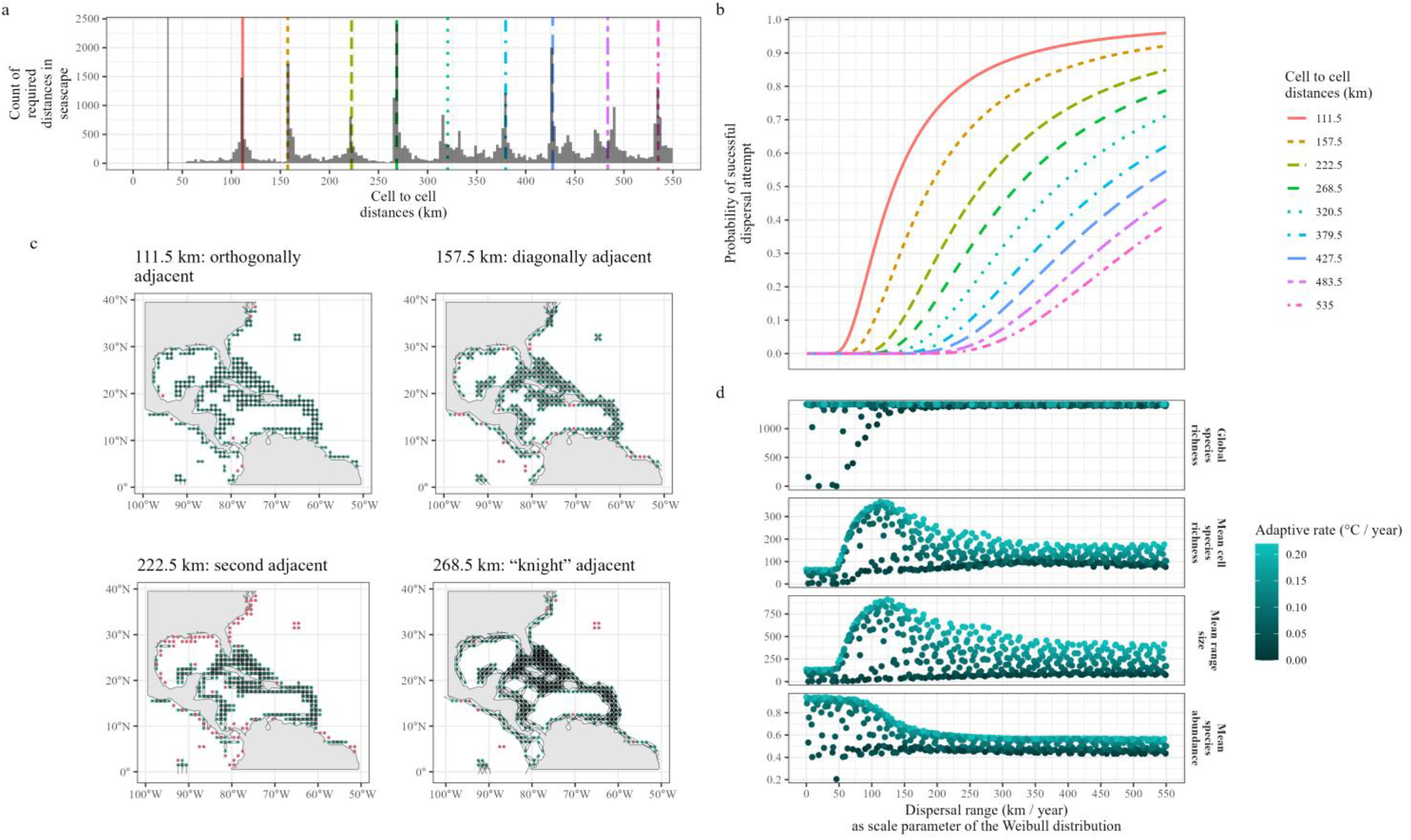
Dispersal-mediated species response thresholds as a function of seascape habitat structure. a) Histogram of cell-to-cell least cost distances in the seascape up to a limit of 550 km corresponding to the maximum species dispersal range parameter across simulations. Coloured, dashed lines mark peaks in this multimodal distribution of distances. The solid vertical grey line is the minimum cell-to-cell distance. b) The probability of species being able to overcome cell-to-cell distance peaks in (a) given their dispersal range parameter. Coloured and dashed lines correspond to the peaks marked in (a). c) Peaks in cell-to-cell distances correspond to adjacency patterns in the 1-degree habitat patch seascape. The panels correspond to the first four peaks in cell-to-cell distances (a). d) Emergent global species responses to cell-to-cell distance thresholds. Inflections in the response curves largely coincide with increases in the probability of overcoming the first few cell-to-cell distance thresholds.

**Supplementary Figure 5:**
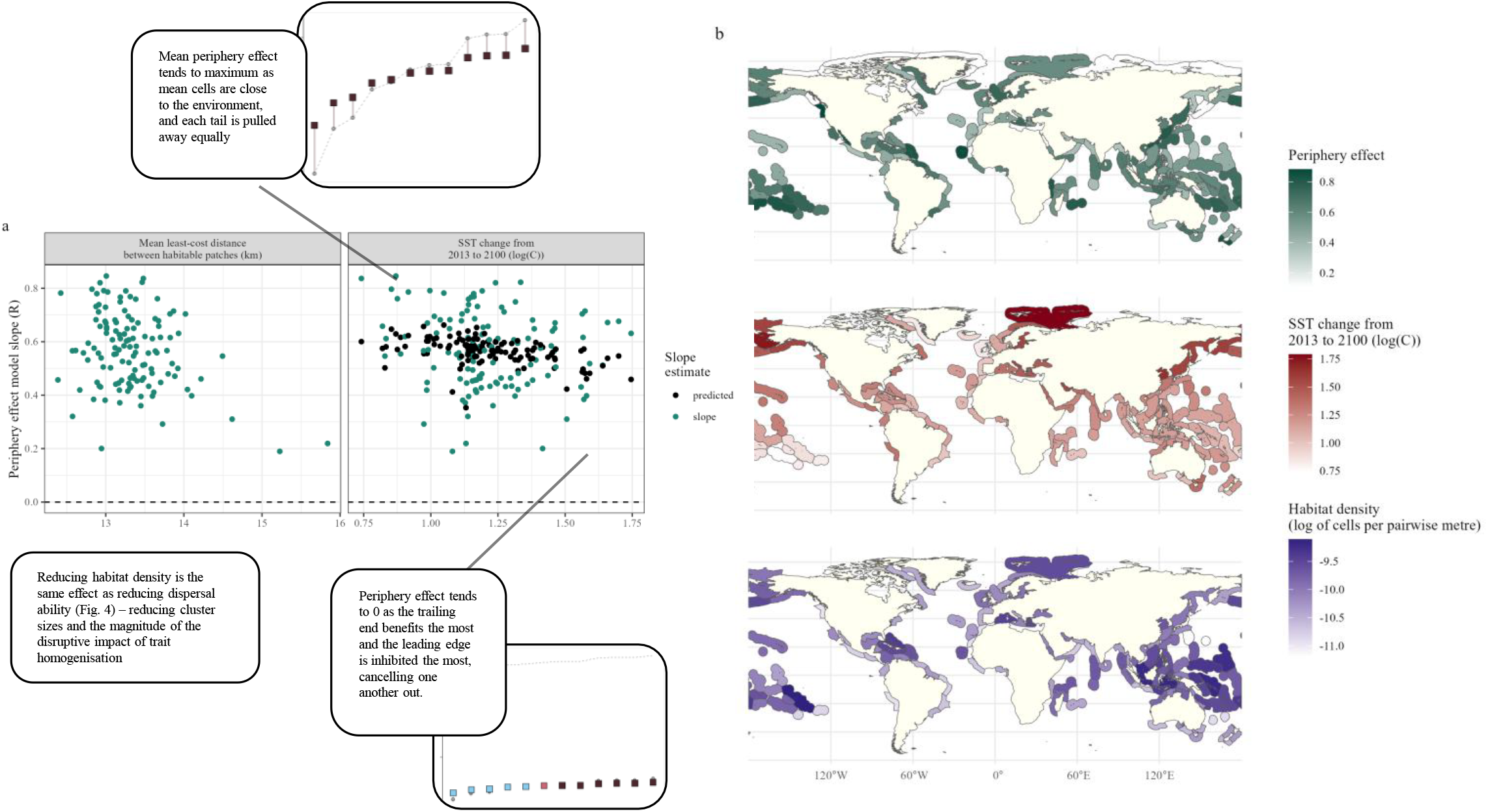
The impact of the seascape (habitat density and thermal change) on the periphery effect. The periphery effect is defined as the relationship between the thermal suitability of a cell and the difference between the local SST and the metapopulation mean SST for a species. a) Each point is the periphery effect model fit across simulations for an ecoregion. Light green points are slope estimates. Dark black points are periphery effect values predicted from ecoregion thermal change values. Only the thermal change values were significant. Boxed panels reflect populations in thermal space following the convention in Fig. 3. b) Maps of ecoregion periphery effects (model slopes), ecoregion thermal change, and ecoregion habitat density. Each ecoregion polygon corresponds to a point in (a).

**Supplementary Figure 6:**
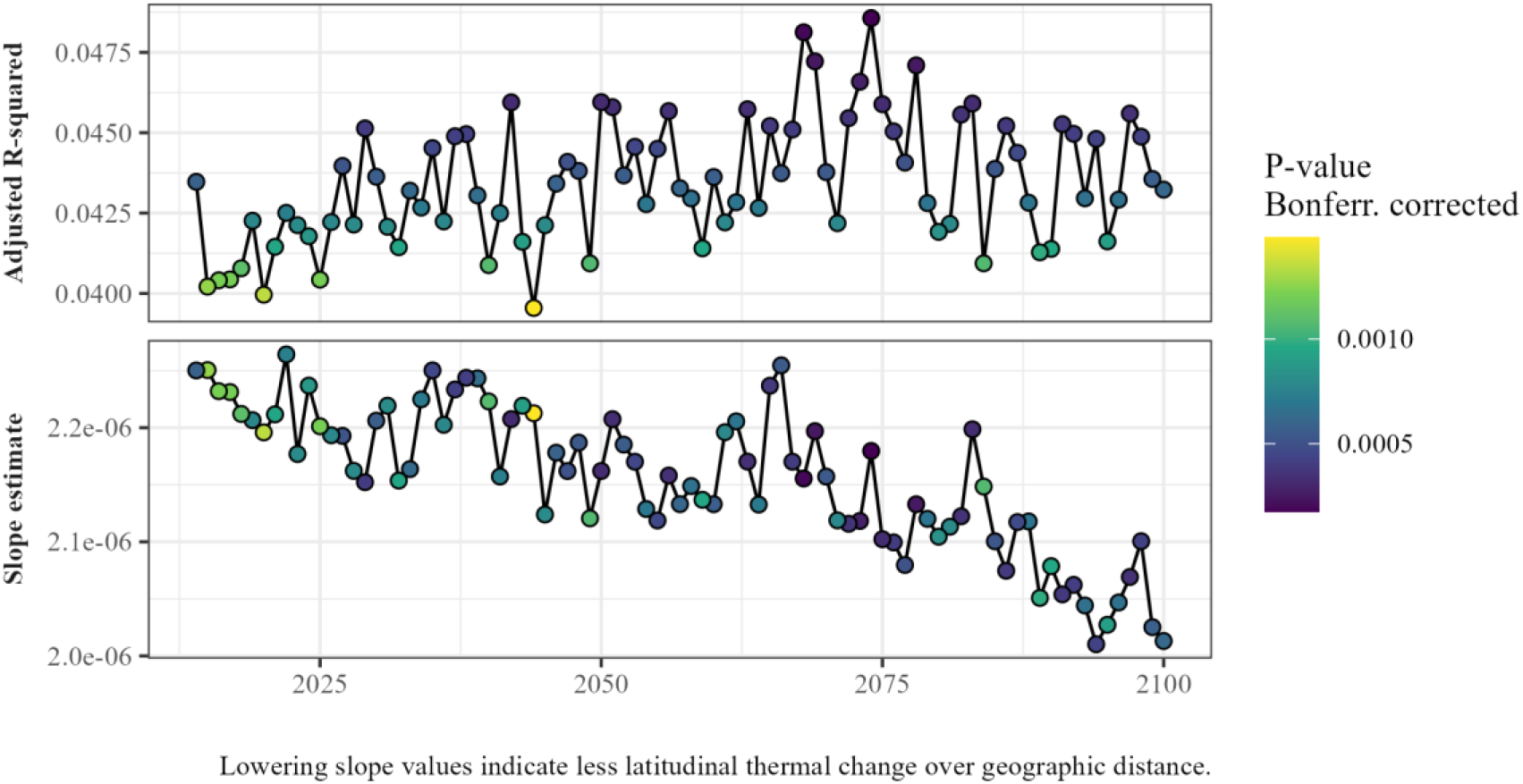
The covariance between geographic least cost distance between habitat cells and the absolute thermal distances in annual mean sea surface temperature between habitat cells, aggregated across latitudes and limited to cells within 550 km of one another – the maximum distance between clustered cells. Higher adjusted R squared values suggest greater variance explanation, lowering slope estimates suggest a less heterogenous thermal seascape over time. Slope estimates and adjusted R-squared values are derived from linear regression fits between the two respective distances value sets.

**Supplementary Figure 7:**
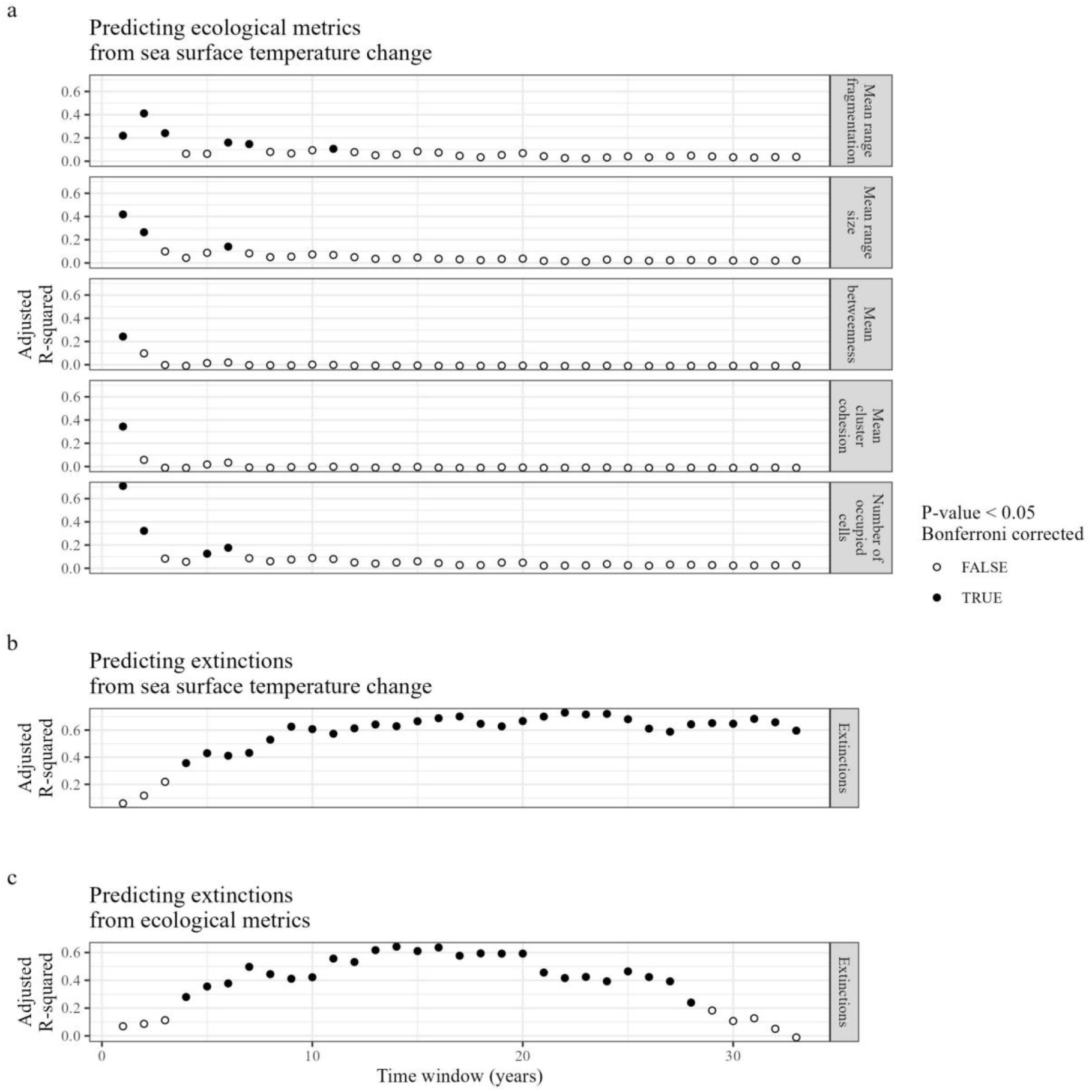
Multistep prediction of species declines and extinctions. Each point is the adjusted R-squared value of a linear model fit. Solid black points indicate a *P*-value < 0.05 after Bonferroni correction for multiple testing. Inversely, white fill points indicate a *P*-value > 0.05. Strip panel labels indicate the response metric. a) Ecological metrics can be predicted using changes in sea surface temperature over the short term. b,c) Extinctions can be predicted from either sea surface temperature change or ecological metrics over the longer term. c) Models fitting ecological metrics to extinctions for time windows above 20 years lose power due to the limited number of time steps in the simulations (86 years).

**Supplementary Figure 9:**
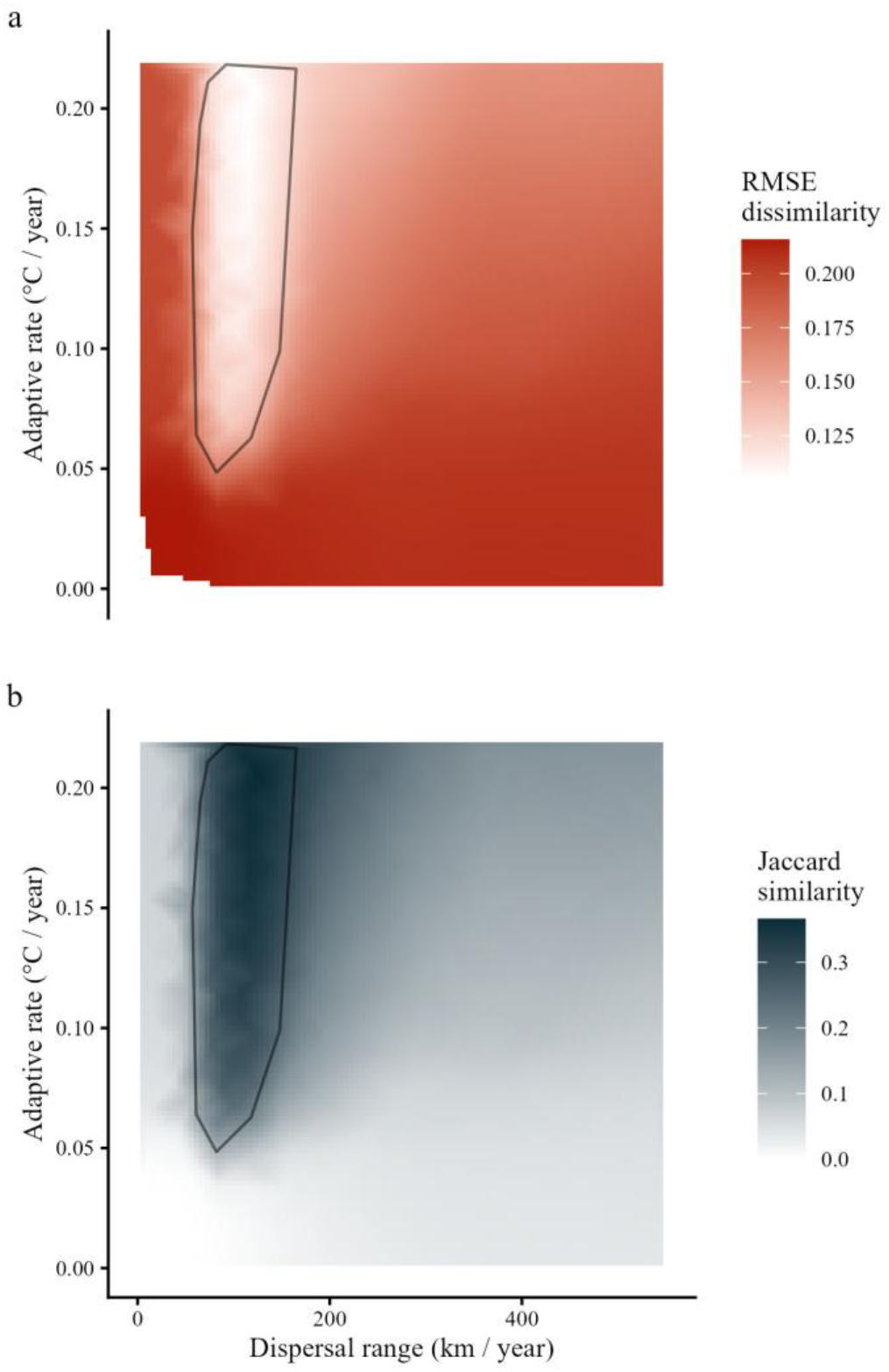
The mean dissimilarity (a) and Jaccard similarity (b) between the species distribution modelling approach by Auber (2019) and the simulation outputs at the year 2100. The grey polygon represents the “Species expansion” response group of simulations (Fig. 2). The mean dissimilarity of species richness patterns was measured using root mean squared error. This does not take into account species identity. Jaccard similarity measure accounts for species composition (temporal turnover).

## Methods

The simulation engine, gen3sis (Hagen et al. 2021), was used to simulate species’ responses to climate change from 2015 to 2100 under different dispersal and adaptive scenarios. To achieve this, we configured simulations whereby species interact with a temporally varying seascape. To explore the impact of adaptation and dispersal, we varied both dispersal ability and thermal adaptive rate across simulations. The configuration of the simulations is detailed below.

## Glossary

*α*: adaptive rate
*a*: abundance
*c*: cluster
*κ*: category
*e*: source cluster
*f*: target cluster
*∈*: Dispersal attempt
*d*: geographic distance
*E*: extinctions
*I*: all cells
*h*: source cell
*i*: focal cell
*j*: target cell
*K*: carrying capacity
*n*: neutral trait
*o*: fragmentation
*λ*: dispersal range
*R*: growth
*ρ*: response variable
*p*: predictor variable
*ϕ*: simulation
*Φ*: all simulations
*s*: species
*S*: all species
*σ*: standard deviation
*T*: sea surface temperature
*t*: time step
*θ*: thermal trait (optimum)
*q*: peripheralness
*Z*: trait value
*μ*: path (in a graph)
*ι*: range, or size
*x*: longitude (1 degree)
*χ*: thermal suitability, or thermal mismatch
*y*: latitude (1 degree)

## Seascape

The gen3sis engine requires two objects describing the seascape: an xy data frame, or table, containing values for each environmental variable for each habitable cell in each time step; and a cell to cell pairwise distance matrix describing the dispersal requirement for migration events between habitable cells. Habitable cells were defined as described in Auber et al. (2022). Briefly, reef locations were defined as within 0.5 degrees of: the Global Self-consistent, Hierarchical, High-resolution Geography dataset (GDHHG) coastline (Wessel and Smith 1996); within the UNEP-WCMC “Global distribution of warm-water coral reefs” (UNEP-WCMC 2010); and covered by SCUBA surveys conducted by the Reef Life Survey (Edgar and Stuart-Smith 2014). This reef habitat covers both tropical and colder waters, excluding Antarctica which was excluded owing to the complications of incorporating ice-sheet dynamics over continental shelf areas. Due to computational limitations, environmental data were aggregated to 1 degree resolution using the aggregate() function in the ‘terra’ package in R (Hijmans 2025). Seascape aggregation resulted in the closure of key geographic passages, so the following manual corrections were made: opening of the Strait of Gibraltar connecting the Atlantic and Mediterranean; opening of the Bab-el-Mandeb Strait connecting the Red Sea and the Arabian Sea; and opening of the Bosphorus connecting the Mediterranean and the Black Sea. The final seascape contained three classes of cell: reef cell, pelagic marine cells, and terrestrial cells – which informed the calculation of the cell-to-cell pairwise distances.

For persistence and dispersal, reef cells were both habitable and passable, pelagic cells were uninhabitable and passable, and terrestrial cells were both uninhabitable and impassable. An exception to this was the Suez Canal: these cells were made 1.5% passable compared to reef and pelagic cells based on documented invasions between the Red Sea and Mediterranean, but remained uninhabitable. Least-cost distances were then calculated between every reef habitat cell using the ‘gdistance’ package in R (Etten 2017). First, a transition object was generated using the reef, pelagic, and terrestrial cell categorisation, using 8 directions, and the “mean” transition function within the transition() function. The resulting transition function was geo-corrected for projected area differences between cells, both longitudinally and latitudinally, then used to calculate least cost paths between habitable reef cells using the costDistance() function.

For each reef cell, SST values were taken from CMIP6 RCP8.5 “business as usual” scenario projections. This includes projecting backward to 1984 and forward to 2100. Projected SST values are derived from the following models: CNRM-ESM2-1, GFDL-ESM4, IPSL-CM6A-LR, MIROC-ES2L, MPI-ESM1-2-LR, NorESM2-MM, UKESM1-0-LL. To assign annual SST values to reef cells, raw monthly model outputs were first spatially reprojected to match the extent and projection of the reef habitat grid. To account for differing coastlines across the climate models, SST values were extrapolated 3 degrees in-land using a sliding window of mean values through the focal() function in the R terra package (Hijmans 2025). These model-specific values were then merged by taking the mean SST value per cell across models for each month. Resulting SST values were then filtered to: remove all non-reef cells, remove all reef cells with no available SST values, and remove cells disconnected from the ocean (an artefact from extrapolating values inland). These filtered monthly SST values were then temporally mean-aggregated to give a single annual SST value per cell from 1984 to 2100.

## Species

Species were seeded into the simulation using estimates of contemporary spatial distributions, local abundances, local thermal optima, and thermal niche ranges. Estimates of spatial distributions were taken from species distribution models compiled in Auber et al. (2022), with occurrence probability values aggregated to 1 degree resolution. Species from these data were filtered down to those identified as “reef-associated” in the Fish Base database (Boettiger, Lang, and Wainwright 2012), and represented in the Fish Tree of Life Project phylogeny (Rabosky et al. 2018). The distributions themselves were restricted to reef cells in the environmental grid and cells with an occurrence probability of less than the species’ median were removed. The remaining probability values were retained as a proxy for initial local abundance at year (Auber et al. 2022).

The thermal niche of each species was simplified to a normal abundance response distribution with a per-cell thermal optimum as the mean, and a species-wide standard deviation. The per-cell species thermal optimum for each occupied cell was calculated as the mean annual SST value for that cell across the years 1981 to 2015. The standard deviation for the thermal niche was calculated as the standard deviation of all annual SST values from 1981 to 2015 across all occupied cells for a species, making it common to all cells in a species.

## Gen3sis function configurations

Our simulations comprise a spatially explicit, population-level metacommunity model with trait evolution, dispersal, and demography across many non-interacting species – built within the gen3sis simulation engine (Hagen et al. 2021). Gen3sis operates on a sequential order of biological processes: dispersal, trait change, and abundance adjustment. This structure abstracts individual-level dynamics, such as mutation, mating, and selection, up to the population level. In this sequence, individuals first colonize new habitable patches via cell-to-cell movement. Following dispersal, the population’s mean trait values shift toward environmental optima. The shift is not modeled explicitly, instead we implicitly consider the combined effect of mutation, mating, genetic mixing and selection. Finally, total cell abundance is recalibrated based on “environmental fit”, reflecting how well the newly adapted trait aligns with local conditions (Waldock et al. 2019). This sequence mirrors the classic “dispersal-mating-selection” paradigm found in low-level simulations but is scaled for higher-level biological organisation. In summary, we execute cell-to-cell dispersal, per cell thermal traits shift to environmental values, per cell thermal traits adjust through metapopulation mixing, then per cell abundance adjusts based on thermal trait and environmental value fit. We configured 500 simulations to run from the year 2014 to 2100 at annual timesteps across the seascape as defined above – made up of habitable reef cells, annual SST values, cell-to-cell distances between reefs, and seeded with our species object. Each simulation differed by two parameter values: dispersal range and adaptive rate, whose values were common to all species within a simulation. We pseudo-randomly varied dispersal capacity and adaptive rate between simulations. Dispersal capacity values for all the species were varied between 0 (no dispersal) and 550 km per time step, derived from tropical fish dispersal estimates (Green et al. 2015). Thermal adaptive rates per time step for all the species were varied from 0 (no adaptation) to 0.22 °C per timestep, which captures the the maximum single-cell temperature difference from one time step to another from 2014 to 2100. The specific values were determined using Sobol sequence values scaled between the range of each parameter, respectively. These parameter values determined the behaviour of the simulations through three core functions: dispersal (movement of species between cells), evolution (change in the thermal optimum trait), and ecology (changes in abundance).

## Dipsersal function

For each species, in each occupied cell, in each timestep, a dispersal attempt is made to every other habitable (reef) cell in the seascape. The distance of the dispersal attempt is sampled from a Weibull distribution with a shape of 2 and a scale set by the dispersal capacity value. If the dispersal attempt distance, *D*, equals or exceeds the corresponding cell-to-cell distance, *d*, in the seascape distance matrix (which is static through time), the dispersal event is successful. The probability of successful dispersal can be expressed with the following survival function,

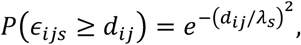

where *i* is the focal cell, *j* is the target cell, *s* is the species, *ε* is the distance of the dispersal attempt, *d* is the geographical distance, and *λ* is the dispersal range.

## Evolution function

### Neutral traits

The neutral trait changes each time step based on a pick from a normal distribution described by,

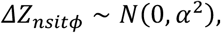

Where *Z*_*nsi*_ is the neutral trait for species *s* in cell *i* at time *t, α*is the adaptive rate, and *ϕ* is the simulation identity.

### Adaptive thermal trait

At each time step, in each species, in each occupied cell, the thermal optimum trait is first moved towards, or to, the local SST value to emulate the population-level consequence of local adaptation. The absolute amount of trait change is the absolute value sampled from a Gaussian distribution with a mean of 0 and a standard deviation value equal to the adaptive rate parameter such that the proposed trait change is given by,

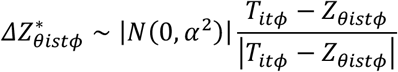

But is limited such that

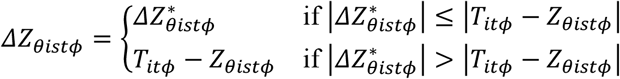

Where *Z*_*θistϕ*_ is the thermal optima for species *s* in cell *i* at time *t* in simulation *ϕ*, and *T*_*itϕ*_ is the mean annual SST value in cell *i* at time *t* in simulation *ϕ*. And where the trait change cannot exceed the difference between the trait value and the environmental value.

### Neutral and adaptive traits

Following its within-cell trait update, both thermal optimum and neutral trait values partially homogenise across geographic clusters of cells. We assume that the effect of trait homogenisation does not scale with geographic distance. Across cells within a cluster, the weighted mean trait value is found – weighted by the abundance value of each cell. Each trait value that has undergone local adaptation is moved towards the abundance-weighted mean by half of the difference (representing equal contribution from implicit random mating) between the weighted mean and the local adaptation value,

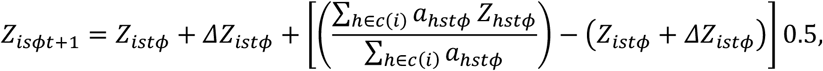

where *Z*_*ist*_ is the trait value for species *s* in cell *i* at time step *t*, and *c*(*i*) is the geographic cluster identity containing cell *i*.

## Ecology

After local adaptation and cluster homogenisation, local abundance values are calculated using a logistic growth function with a both the growth rate and carrying capacity scaled by the thermal suitability. Thermal suitability, and therefore both the carrying capacity and growth rate, are defined by the Gaussian decay from the local SST to the thermal optimum. *I*.*e*., the realised distribution density over the optimal distribution density,

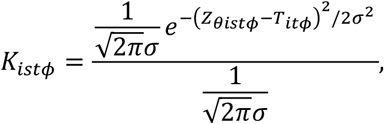

Where *K*_*ist*_ is the carrying capacity for species *s* in cell *i* at time *t*, and *σ* is the Gaussian standard deviation. The annual growth is then calculated as,

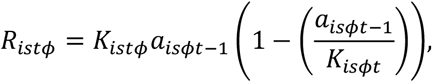

Where *R*_*ist*_ is the growth for species *s* in cell *i* at time *t*. The final abundance value is then hard-limited to the local carrying capacity, and driven to extinction below abundance values of 0.1,

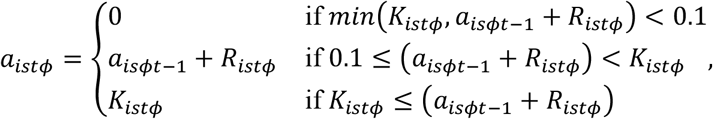

Where *a*_*ist*_ is the abundance for species *s* in cell *i* at time *t*.

## Response metric definitions

Output metrics were calculated for every year for each simulation run across three levels of aggregation. The levels of aggregation are: per cell, metrics that capture the properties of each cell in the seascape to allow spatial information; per species, metrics that are aggregated across a species to allow inter-species comparisons; and per simulation, global metrics that are aggregated across the entire simulation for that timestep for broad inter-simulation comparisons. *I*.*e*.,

- Simulation - single metric value per year, per simulation.
- Species - single metric value per species, per year, per simulation.
- Cell - single metric value per cell, per year, per simulation.

All metrics are time dependent and per simulation.

## Cell metrics

### Cell richness

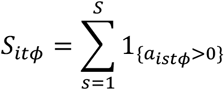

### Total cell abundance

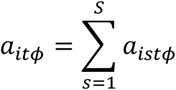

### Mean cell abundance

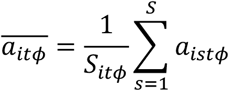

### Mean thermal optimum

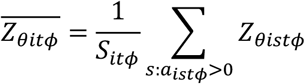

### Mean cell tolerance mismatch

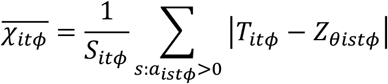

### Betweenness centrality

Betweenness was calculated for each geographic cluster within a species using the `betweenness()` function in the ‘igraph’ package in R, following the methodology of Freeman (1978). From these per-cluster betweenness values, the minimum, mean, and maximum were calculated per species, per time step (year).

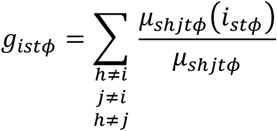

where *h* is a source cell, *j* is a sink cell, *i* is the cell of interest, and *s* is the species.

#### Mean betweenness

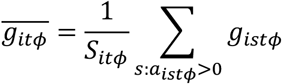

#### Minimum betweenness

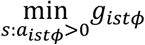

#### Maximum betweenness

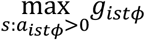

### Phylogenetic diversity

Phylogenetic diversity per cell as calculated by `picante::pd()` (Kembel et al. 2010) using all species present in that cell and the fish tree of life phylogeny (Rabosky et al. 2018).

### Mean pairwise distance

Phylogenetic mean pairwise distance between species in a cell was calculated using `picante::mpd()` (Kembel et al. 2010) and all species present in that cell (Rabosky et al. 2018).

### Variation in pairwise distance

Variation in mean pairwise distances between species in a cell *S*_*sitϕ*_. This was calculated as the variance of the distance matrix is derived from the fish tree of life phylogeny (Rabosky et al. 2018; Chang et al. 2019) using the `stats::var()` function (R Core Team 2022).

## Species metrics

### Total species abundance

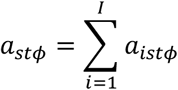

### Mean species abundance

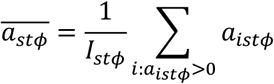

### Mean species thermal optimum

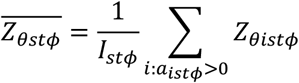

### Standard deviation of species thermal optima

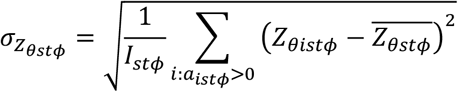

### Range size

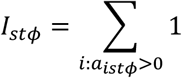

### Mean species tolerance mismatch

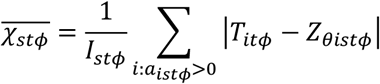

### Range of thermal optima

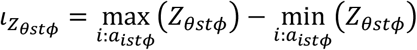

### Cluster count

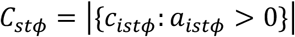

### Cluster size

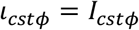

### Mean cluster size

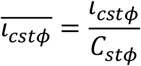

### Maximum cluster size

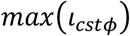

### Minimum cluster size

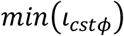

### Mean inter cluster distances (km)

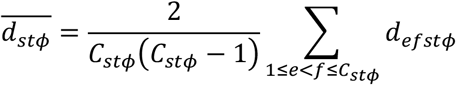

### Cohesion

Cohesion was calculated for each geographic cluster within a species using the `cohesion()` function in the ‘igraph’ package in R (Csárdi et al. 2025), following the methodology of White and Harary (2001). From these per-cluster cohesion values, the minimum, mean, and maximum were calculated per species, per time step (year).

### Betweenness

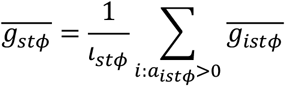

### Fragmentation

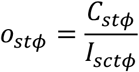

## Simulation metrics

### Global species richness

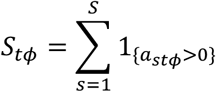

### Occupied cells

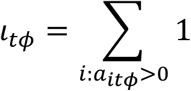

### Mean species richness per cell

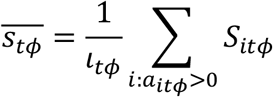

### Total abundance across all species

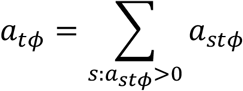

### Mean total abundance

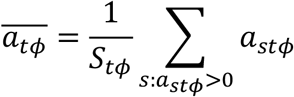

### Mean range size of species

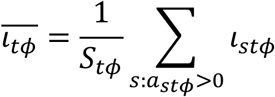

### Mean cluster size of species

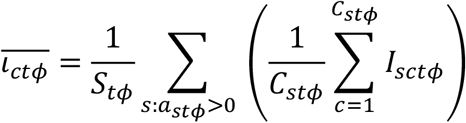

### Mean number of clusters per species

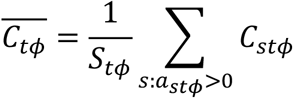

### Mean of the mean thermal optimum of species

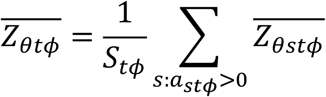

### Mean tolerance mismatch

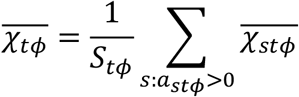

### Mean/min/max distance between clusters

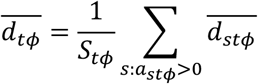

### Mean/min/max betweenness

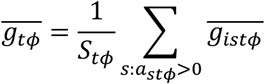

### Phylogenetic diversity

Calculated using the pd() function in the R picante package (Kembel et al. 2010) on all surviving species in the simulation.

### Mean pairwise phylogenetic distance

Calculated using the mpd() function in the R picante package (Kembel et al. 2010) on all surviving species in the simulation.

### Variation in pairwise phylogenetic distances

Calculated using the base R var() function (R Core Team 2022) on all surviving species in the simulation.

### Mean range fragmentation

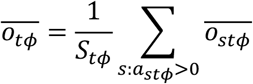

## Unsupervised simulation clustering

Simulations were clustered into response groups in continuous space using a principle components analysis, and into discrete groups using the kmeans algorithm.

Clustering was based on the simulation metrics of mean range size, global species richness, mean species richness per cell, mean abundance per cell, the number of occupied cells, metapopulation fragmentation, and the mean distances between metapopulation clusters. Mean range fragmentation, mean range size, and mean species richness per cell were all log transformed to better fit the assumption of normality for the PCA. Global species richness was heavily right-skewed, so this was recalculated as “extinctions” by subtracting species richness values from the initial number of species in the simulation. This was then log transformed. Giving,

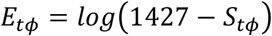

where 1427 is the number of species at simulation initialisation. Metric values were then mean-centred and scaled by their standard deviations using the `scale()` function in R’s `stats` package (R Core Team 2022). The PCA was carried out on the centred and scaled metrics using the `prcomp()` function with default arguments.

Before kmeans clustering, the most suitable value for the number of groups, k, was determined by maximising the the average silhouette width for each simulation in response space using the `fviz_nbclust()` function from the `factoextra` package in R (Kassambara and Mundt 2020). Five groups returned the maximum width with a value of 0.49. Simulations in each group roughly corresponded to distinct responses areas in parameter space, and were labelled accordingly. Five simulations experienced global extinction of all species by the year 2100, making it impossible to generate any response metrics. These were manually binned into a sixth group for “global extinction”.

## Latitudinal Diversity Gradient

Across all simulations within each group, the mean species richness per cell at each degree of latitude was calculated for each year in a simulation as,

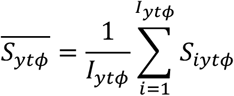

where *S*_*ytϕ*_ is the species richness at latitude *y* at time *t* in simulation *ϕ*, and *S*_*iytϕ*_ is the species richness at cell *i* at latitude *y* at time *t* in simulation *ϕ*. The mean per response group was then calculated as,

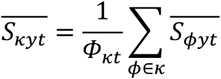

where *κ* is the response group. These values give a time series of mean richness across latitude for each of the five extant response scenarios.

## Defining the LDG response

For each simulation, at each year, in each hemisphere, the latitudinal diversity gradient was defined using both a uniform and Gaussian model fit using functions from R’s ‘stats’ package (R Core Team 2022). To fit and estimate the parameters of the Gaussian model to the latitudinal richness distribution, the `nls()` function was used.

We initialised the Gaussian parameter (mean latitudinal species richness, latitude of peak species richness, and the standard deviation of latitudinal species richness) search using: the Gaussian function; species richness mean and peak species richness values from a LOESS fit using the `loess()` function with a latitudinal span of 0.5; and standard deviation values calculated directly from mean latitudinal species richness values using the `sd()` function. The uniform model was fit using the `lm()` function where the mean latitudinal richness is defined by the constant, 1. Uniform and Gaussian model fits were then compared using AIC scores as calculated using the `AIC()` function and the best fitting model for each simulation, year, and hemisphere was retained. For instances where the Gaussian parameter fitting did not converge, the uniform model was assumed to be a better fit. *P*-values for all retained models were Bonferroni corrected for multiple testing.

## Temporal Turnover

Turnover in species between the years 2014 and 2100 was characterised in each cell in each simulation by colonisations, extinctions, species richness turnover, and Jaccard’s species composition similarity. Colonisations were defined as the number of species present in a cell at 2100 that were not present in 2014. Extinctions were defined as the number of species not present in a cell at 2100 that were present in 2014. Species richness turnover was defined as the species richness at 2100 minus the species richness at 2014. Jaccard’s species compositional similarity was calculated as the number of species present at both 2100 and 2014 over the combined (union of the) species pool across 2014 and 2100. These metrics were then further mean aggregated across latitudinal bands and dispersal/adaptive categories, as was done for species richness.

## Temporal turnover vs LDG response

The covariation in temporal turnover in species composition and the shift in latitude and standard deviation of the LDG was characterised using linear regression fits. For both LDG latitude and latitudinal standard deviation in each hemisphere, a simple linear regression was fit using the mean Jaccard’s similarity in species composition between 2014 and 2100 for each simulation as a predictor variable.

## Gene flow and metapopulation adaptive dynamics sub-model

Figure 3.

To highlight the impact of dispersal mediated gene-flow on the adaptive dynamics of metapopulations, outputs from the year 2025 from two simulations (run IDs: 362 and 328) with comparable adaptive rates, but with high and low respective dispersal ranges were isolated from the main batch of simulation runs. From each of these simulation outputs we extracted the species information for *Macrophayngodon bipartitus* from within the Western Indian Ocean. To illustrate trait homogenisation dynamics across a seascape variable in both thermal conditions and connectivity in our model, we manually extracted occupied cells around the Scattered Islands, Madagascar, Mauritius, Reunion, and Vingt Cinq (cell IDs: 36584, 36944, 36945, 36946, 35867, 39104, 38744, 38384, 38026, 36949, 36590, 38395). For the high dispersal scenario (run ID 362), all occupied cells were grouped into a single metapopulation cluster. For the low dispersal scenario (run ID 328), cells were split into 6 metapopulation clusters as determined by their dispersal range of 118 km. Sea surface temperature values for occupied cells were pulled from the seascape object directly. To focus on the trait homogenisation and adaptive dynamics, all the the species’ thermal optima were standardised across cells by manually setting them to be the local sea surface temperature value minus the mean thermal change per year across the seascape object. The evolution function as described in the configuration section was then applied to the occupied cells using the dispersal range and adaptive rate values from the two chosen scenarios.

The consequence of trait homogenisation across metapopulation clusters on per cell adaptive success was categorised based on their values relative to both their starting values, and their values after adaptation alone (before the trait homogenisation function is applied). If the final thermal optimum value had a closer fitness than both the starting and post-adaptation values then trait homogenisation “increases fitness”. If there was only one cell in a cluster and did not dispersal between cells then there was “no trait homogenisation”. If the final value had a worse environmental fitness to the adaptation-only values, then it “decreases fitness”, and it “causes maladaptation” if the final value was less fit than the starting value.

## Extinction dynamics

Figure 5.

Expectations for the survivability of species through time were derived from assessing occupied cell habitability per time step. For all species, cells across their entire range were derived from their starting distributions, as well as their per-cell thermal optima and standard deviations. These values, for both traits and cell occupancy, were held static through time, whist the annual mean sea surface temperature was allowed to increase according to the seascape input object. For each species, in each cell, in each time step, the ecology function (which determines per cell abundance and extinctions) was run using a best case scenario initial abundance (1) to determine abundance change and possible local extinction. The output of these calculations show, given no dispersal, thermal adaptation, and maximum starting abundance, if a cell would be habitable at each year given the projected thermal regime. From these values, global extinction of a species per time step was assigned if all per cell abundance values were driven to 0. The first year where a species experiences global extinction was recorded as the time at which that species’ range was no longer habitable.

## Dispersal thresholds

Supplementary Figure 4.

Response metrics to dispersal ability showed non-linear responses. To understand the points of inflection in these non-linear responses, the seascape was characterised through pairwise cell-to-cell distances. The dispersal parameter space has a maximum value of 550 km, so cell-to-cell distances greater than 550 km were removed as response values beyond the parameter space limits are not comparable with parameter values. Visually, the distribution of the pairwise distances between the remaining cells contained a series of highly distinct peaks (Supplementary Fig. 4a). To characterise these peaks, distances were categorised into 500 bins (1.1 km bins) between 0 and 550 km to cover the entire dispersal ability space. Using the midpoint of the distance bins as a predictor variable and the count of observations per bin as a response variable, a LOESS model was fitted with a span of 0.1 using the `stats` package in R (R Core Team 2022). This produced a multi-peaked curve from which local maxima were derived using the the `find_peaks()` function in the `ggpmisc` R package, using a span of 3 (Aphalo, Slowikowski, and Mouksassi 2025). To remove noisy local maxima, local maxima with a count of less than 100 were removed. Probabilities for species to overcome each peak in cell-to-cell distances was calculated for dispersal ranges from 0 to 550 km. This was done using the `pweibull()` function from the `stats` package in R (R Core Team 2022), using the same shape value as used in the gen3sis dispersal function.

## Comparison of species distribution model outputs and gen3sis simulations

Supplementary Figure 8.

Direct comparison of distribution patterns derived from the gen3sis simulations were made with species distribution modelling (SDM) outputs by Auber et al. (2022) for the year 2100. Outputs were first wrangled to be made comparable. SDM species probability values were aggregated from a resolution of 0.25 degrees to 1 degree by finding the mean value per 1 degree cells across its constituent 0.25 degree cells. These probability values were then converted into binary presence/absence records by recording any probability greater than 0 to be 1 and converted into a presence/absence matrix using in-house R scripts. Similarly, gen3sis outputs were converted into presence/absence matrices using in-house R scripts to match the format of the SDM matrix. Species richness per cell for both the SDM outputs and the gen3sis simulation outputs were calculated by summing the number of species present in each cell. Per cell richness values were then compared using the root mean squared error difference, and Jaccard’s similarity was calculated using in-house R functions.

## Metapopulation periphery effect

Supplementary Figure 5.

The dependence of the impact of gene flow on metapopulation cells on their distance from metapopulation mean thermal trait values and environmental temperature, was defined statistically. To allow a comparison of seascape structure on this peripheral effect, cells were split into ecoregions (Spalding et al. 2007).

For simulation outputs at the year 2100, the thermal optimum per species, per cell, was extracted. Cells that were part of a metapopulation containing fewer than 3 cells were removed. The thermal suitability of the remaining cells across all species was calculated as the absolute difference between the thermal optimum trait value and the local sea surface temperature. Per cell, cluster, species, and simulation, the “environmental peripheralness” was calculated as the absolute difference between the SST value of that cell, and the mean SST value of that species’ metapopulation cluster, defined as:

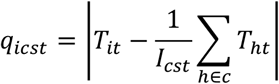

Due to computational limitations, only a random sample of 50,000 cells across species were retained per simulation, and 20,000 retained per ecoregion across simulations for model fitting. Ecoregions subsequently containing fewer than 5 observations were removed. This resulted in 136 retained ecoregions.

The relationship between thermal suitability (the difference between species’ thermal trait optima and local environmental temperature) and how far a cell was from the environmental metapopulation mean was defined using a phylogenetic mixed-effects model using generalised least squares. Thermal suitability was the response variable, environmental peripheralness per ecoregion was the predictor variable, simulation identity was a grouping factor, and a Brownian phylogenetic covariance structure accounted for shared evolutionary history amongst species (Rabosky et al. 2018). This was implemented using `pglmm()` function from the `phyr` package in R.

The relationship between the slopes of these periphery effect models was then characterised by a linear fit to seascape characteristics. First, *P*-values across periphery effect model fits were Bonferroni corrected and model fits with *P*-values above 0.05 were removed. Seascape characteristics were characterised as the mean change in sea surface temperature from 2014 to 2100 for each ecoregion, and the a measure of habitat density - the number of habitable cells over the mean pairwise distances between those cells. Each of these measures was then log-transformed to satisfy the linear regression assumption of normality. The significant slope values describing the strength of the periphery effect for each ecoregion were then fitted as a response variable to mean cell-to-cell distances within an ecoregion and mean thermal change between 2014 and 2100 per ecoregion in a simple linear regression using `lm()` in R’s `stats` pacakage (R Core Team 2022). These predictor variables were then selected stepwise for best model fit using the `step()` function.

## Pairwise thermal and geographical distance correlations

Supplementary Figure 6.

The correlation between pairwise sea surface temperature differences and pairwise cell-to-cell distances was evaluated for each time step in the simulations. Pairwise cell-to-cell distances were taken directly from the distance matrices used as input for the simulations. Pairwise thermal distances between all habitable cells at all time steps were calculated as the absolute differences between sea surface temperature values between all cell pairs.

For each time step, a simple linear regression was fit for each time step with all pairwise distances in sea surface temperatures between cells as the response variable, and geographic distances as the predictor variable. *P*-values were then Bonferroni corrected for multiple testing.

### Species response and extinction time series

Supplementary Figure 7.

To understand global levels of species decline or expansion, or extinction, we built three models:

1. predicting global extinctions per time step across simulations using change in sea surface temperature;
2. predicting species response metrics using change in sea surface temperature;
3. predicting species extinctions using species response metrics.

Species response metrics included: mean species abundance, mean species range fragmentation, mean species range size, mean species metapopulation cohesion, mean species metapopulation betweenness, and mean species inter-cluster distances. For all three of these models, the response variables were the mean changes in response metric values across all simulations from the target year from the previous year,

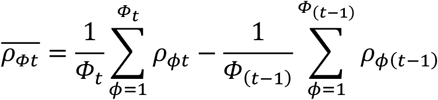

where *ρ* is the response variable, *Φ* is the simulation, and *t* is the time. Since our simulations carry over the simulation state from one year to to the next in a contiguous time series, we factored in multi-year cumulative responses to thermal change. We did this by building each model across a range of time windows, *i*.*e*., the change in predictor value over a single time step, two time steps… up to 40 time steps, such that,

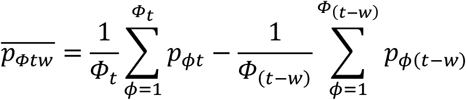

where *p* is the predictor variable and the time window values belong to the set,

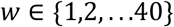

Three sets of models were then built to predict 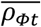 using 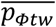 for each predictor, response, and time window value, *w*. Models were fit using the `lm()` function from R’s `stats` package (R Core Team 2022). For models predicting extinctions per year using sea surface temperature change as a predictor variable, we found that the response contained thresholds, or tipping points, in the magnitude of change through time whereby after a certain thermal change value, the extinctions no longer increased. We therefore fitted segmented linear regressions to each time window combination using the `segmented()` function from R’s `segmented` package (Fasola, Muggeo, and Küchenhoff 2018). All model *P*-values were Bonferroni corrected for multiple testing by the number of time windows (40) per predictor-response combination.

